# Aquatic long-term persistence of *Francisella tularensis* ssp. *holarctica* is driven by water temperature and transition to a viable but non-culturable state

**DOI:** 10.1101/2022.02.18.480867

**Authors:** Camille D. Brunet, Julien Peyroux, Léa Pondérand, Stéphanie Bouillot, Thomas Girard, Éric Faudry, Max Maurin, Yvan Caspar

## Abstract

*Francisella tularensis* is a highly virulent bacterium causing tularemia zoonosis. An increasing proportion of infections occur through contaminated hydro-telluric sources, especially for the subspecies *holarctica* (*Fth*). Although this bacterium has been detected in several aquatic environments, the mechanisms of its long-term persistence in water are not yet elucidated. We evaluated the culturability and the viability of a virulent *Fth* strain in independent microcosms filled with nutrient-poor water. At 37°C, the bacteria remained culturable for only one week, while culturability was extended to 6 weeks at 18°C and up to 11 weeks at 4°C. However, while the viability of the bacteria declined similarly to culturability at 37°C, the viability of the bacteria remained stable overtime at 18°C and 4°C for more than 24 months, long after loss of culturability. We identified water temperature as one of the major factors driving the aquatic survival of *Fth* through a transition of the whole *Fth* population in a viable but non-culturable (VBNC) state. Low temperature of water (≤18°C) favors the persistence of the bacteria in a VBNC state, while a temperature above 30°C kills culturable and VBNC *Fth* bacteria. These findings provide new insights into the environmental cycle of *Francisella tularensis* that suggest that the yet unidentified primary reservoir of the subspecies *holarctica* may be the aquatic environment itself in which the bacteria could persist for months or years without the need for a host.

## Introduction

*Francisella tularensis* is a Gram-negative bacterium causing the zoonosis tularemia. It is a highly virulent human pathogen classified in category A of potential agents of biological threat by the US Centers for Disease Control and Prevention [1]. Two subspecies are associated with human tularemia: *F. tularensis* ssp. *tularensis* (*Ftt*) (type A strains), only present in North America; and *F. tularensis* ssp. *holarctica* (*Fth*) (type B strains), spread all over the Northern Hemisphere, with a few strains identified in the last decade in Australia [1,2].

Terrestrial and aquatic lifecycles of *F. tularensis* have been described but remain not fully characterized despite many decades of research [3]. Especially, the survival of the bacteria in hydro-telluric environments is still under active investigation [4,5]. The terrestrial animal reservoir of *F. tularensis* is large, but lagomorphs and small rodents are considered primary sources of human infections. Recent data corroborate that the aquatic lifecycle of the subspecies *Fth* may be predominant over the terrestrial lifecycle, in particular for the persistence of the disease in the environment, as initially suggested by Jellison [5,6]. This aquatic cycle involves mainly mosquitoes, mosquito larvae, and aquatic rodents [3]. In Northern Europe, mosquitos can transmit *Fth* after larva contamination in water and consequently be responsible for large outbreaks [7–10]. Cases of tularemia related to water have also been described after an aquatic activity (e.g., swimming or canyoning) [11,12] or through drinking or using contaminated water [13,14].

Some studies suggested the potential persistence of this bacterium in aquatic environments over long periods. Genomic studies have confirmed that diverse clones of *Fth* survive for a prolonged period and that a single clone may be responsible for human or animal cases of tularemia over several decades (up to 70 years) [15–17]. Multiple independent respiratory infections with *Fth* strains acquired from the environment over a short period were observed during an outbreak in Sweden in 2010 and in France in 2018, arguing in favor of environmental changes acting as the trigger of these outbreaks [17,18]. Analysis of exposition factors suggested environmental contamination, presumably through aerosols originating from an unidentified environmental reservoir [5]. Low temperature and salinity have been identified to impact the duration of culturability of *Fth*. It has been described that this bacterium can remain culturable up to 70 days at 8°C [19,20], ten days in fresh water at room temperature, 21 days in seawater, and 45 days in brackish water [21]. Recently while studying biofilm formation of *F. tularensis* in aquatic environments, Golovliov *et al*. identified that *Fth* remained culturable and infectious in a mice model after 24 weeks of incubation at 4°C in low nutrient water containing 9 g/L of NaCl. They suggested that this improved survival at low temperature in freshwater may be a critical mechanism to help the bacteria overwinter and survive between host-associated replication events [4]. In such situations the bacteria may choose to switch to a dormancy state that reduces competition with actively growing cells. Among potential persistence and/or quiescence mechanisms identified in bacteria, bacterial switch to a viable but non-culturable (VBNC) state that has been poorly studied in virulent *F. tularensis* strains [20,22]. Initially described in 1982 for *Escherichia coli* and *Vibrio cholerae* [23], the VBNC state corresponds to bacteria that lose their ability to grow, may change their shape and lose their virulent traits, although remaining still alive. The VBNC state is induced during a stress such as nutrient starvation, physicochemical changes of the environment, or thermal shock. It has already been identified that the virulence-attenuated live vaccine strain (LVS) of *Fth* is able to survive in a VBNC state at least 140 days at 8°C [20]. Survival of a fluorescent *Fth* strain in a VBNC state up to 38 days has also been described in the control conditions of a co-culture experiment with protozoan using a gfp-modified *Fth* strain [22].

Consequently, our goal was to investigate the role of water temperature and salinity on the persistence of a virulent human strain of *Fth* in water and explore the possibility of a transition of *Fth* into a VBNC state triggered by these factors.

## Material and methods

### Bacterial strains and preparation of aquatic microcosms

All culture assays were performed in a BSL3 laboratory. We used the fully virulent *Fth* biovar I clinical strain CHUGA-Ft6 (genome accession: VJBK00000000) [15]. This strain was grown on Polyvitex-enriched chocolate agar plates (PVX, BioMérieux, Marcy l’Etoile, France) incubated at 37°C in a 5% CO2-enriched atmosphere. The *F. tularensis* collection of French National Reference Center for *Francisella* is approved by the Agence Nationale de Sécurité du Médicament et des produits de santé (France) (ANSM, authorization number ADE-103892019-7).

Six independent aquatic microcosms were defined, consisting of 6 aliquots of the same environmental water sample from the Rhône-Alpes region in France (send for analysis in the water laboratory Abiolab-Asposan, Monbonnot-Saint-Martin, France; Table S1). Microcosms were incubated at 4°C, 18°C, and 37°C, and supplemented with either 0 or 10 g/L of NaCl. Each condition was tested in biological triplicate. Bacterial suspensions were prepared in PBS and adjusted to 10^9^ CFU.ml^-1^, and 25 mL were added to 225 mL of environmental water previously sterilized using a 0,22μm filter.

### Monitoring of culturability and viability of bacteria in water

The culturability and viability of bacteria in the six environmental models were monitored each week. The culturability was measured by CFU counts after plating 100μL of serials dilutions of each microcosm and on PVX agar plates after 48h incubation at 37°C. The viability of the bacteria was determined using qPCR after PMAxx™ Dye treatment (Biotium San Francisco, US) allowing specific DNA amplification of viable bacteria only. In brief, 1 mL of bacteria suspension was added to 250 μL of enhancer for Gram-negative Bacteria (Biotium San Francisco, US). PMAxx™ Dye was dissolved in H_2_O at 5mM and added to a bacterial solution at a final concentration of 25μM. After 10 min of incubation in the dark, samples were exposed 30 min to light with GloPlate™ Blue LED Illuminator (Biotum, San Francisco, US). Bacterial-PMA suspensions were centrifugated at 11,000g for 10 min, and DNA was extracted using NucleoSpin Blood Kit (Macherey Nagel, Hoerdt, France) according to manufacturers’ recommendations. At each sampling point, DNA extraction of 1 mL of bacterial suspension without PMA treatment was realized in parallel to determine amplification of total DNA present in the samples. At each sampling point, control with dead bacteria for PMAxx™ Dye was realized using a 1 mL suspension of bacteria previously lysed. Each qPCR reaction contained 10μL of Master Mix EvaGreen 2X (Biotium, San Francisco, US), 1μM of each primer, 5μL of DNA template, and 1μL of sterile H_2_O. The 23S ribosomal RNA gene was amplified using the following forward (5’-CATACGAACAAGTAGGACGG-3’) and reverse (5’-GCAAGCGGTTTCAGATTCTA-3’). The qPCR was performed using a LightCycler 480 instrument (Roche, Meylan, France) and SYBRGreen channel, with the following protocol: initial denaturation at 95°C for 5 min, followed by 40 cycles of 95°C for 5s and 60°C for 30s. Melting curve analysis was performed from 57°C to 99°C. A negative control (H_2_O) was included in each qPCR run. The viability was evaluated by the Cycle threshold (Ct) of DNA amplification of living Bacteria. Statistical analyses were performed by student t-test.

To compare the shape and the length of VBNC and culturable bacteria, pictures of bacteria incubated in water at 18°C without NaCl one hour for culturable bacteria, and 6 months for VBNC bacteria labeled with Syto9 were analyzed by Image J software and Microbe J plugging [24]. Bacterial morphology was described by parameters: area (0.1-1.2μm); length (0.2-1.4 μm); width (0-1.4 μm); circularity (0.3-max μm). Parameters were calculated for 1055 bacteria in each sample.

### Bacterial viability of VBNC cells after temperature change of microcosm

Several months after the loss of culturability of *Fth* in water, 5 mL of microcosm at 4°C were transferred at 18°C, 30°C and 37°C. 5 mL of microcosm at 18°C were transferred at 4°C, 30°C and 37°C. After 7 and 14 days, the viability was evaluated by qPCR after PMAxx™ Dye treatment. Statistical analyses were performed using R (version 4.0.3) for the comparison of multiple groups by one-way ANOVA. False discovery rate (FDR) correction was applied for pairwise t-tests.

### Biofilm quantification

Biofilm quantification was performed on the bacterial suspensions evolved in nutrient-poor water in T75 culture flasks, one year at 4°C, six months at 18°C, or four months at 37°C after inoculation. Negative control consisted in fresh culturable bacteria incubated for one hour in water. Each condition was tested in a biological duplicate. After incubation, the culture medium was aspirated, and flasks were washed three times with PBS. 5 mL of Crystal violet (0,2% w/v) were added, and flasks were re-incubated for 15 minutes. Crystal violet was washed three times with PBS, and biofilm was solubilized by 1 mL of ethanol 95%. 200μL of this biofilm was added to microtiter plates in three wells, and biofilm was quantified by measuring absorbance at 570nm. Statistical analyses were performed by student t-test.

## Results

### Culturability of F. tularensis ssp. holarctica extends to 11 weeks at low temperature

In low nutrient-containing water, at 37°C, the culturability of the virulent clinical strain of *Fth* biovar I decreased from 10^8^ to 0 CFU/mL in 8 days (Figure 1a). However, the culturability of bacteria extended dramatically when reducing the temperature of water microcosms. At 18°C, bacteria decreased from 10^8^ to 0 CFU/mL in 6 weeks (Figure 1b). At 4°C, culturability of bacteria declined even more slowly with a complete absence of growing colonies only 11 weeks after inoculation of the water sample (Figure 1c). At 37°C, NaCl concentration enrichment of the microcosm at 10g/L conferred only a slight transient survival advantage to the bacteria (Figure 1a). No significant differences were observed at 18°C or 4°C.

**Figure 1:**
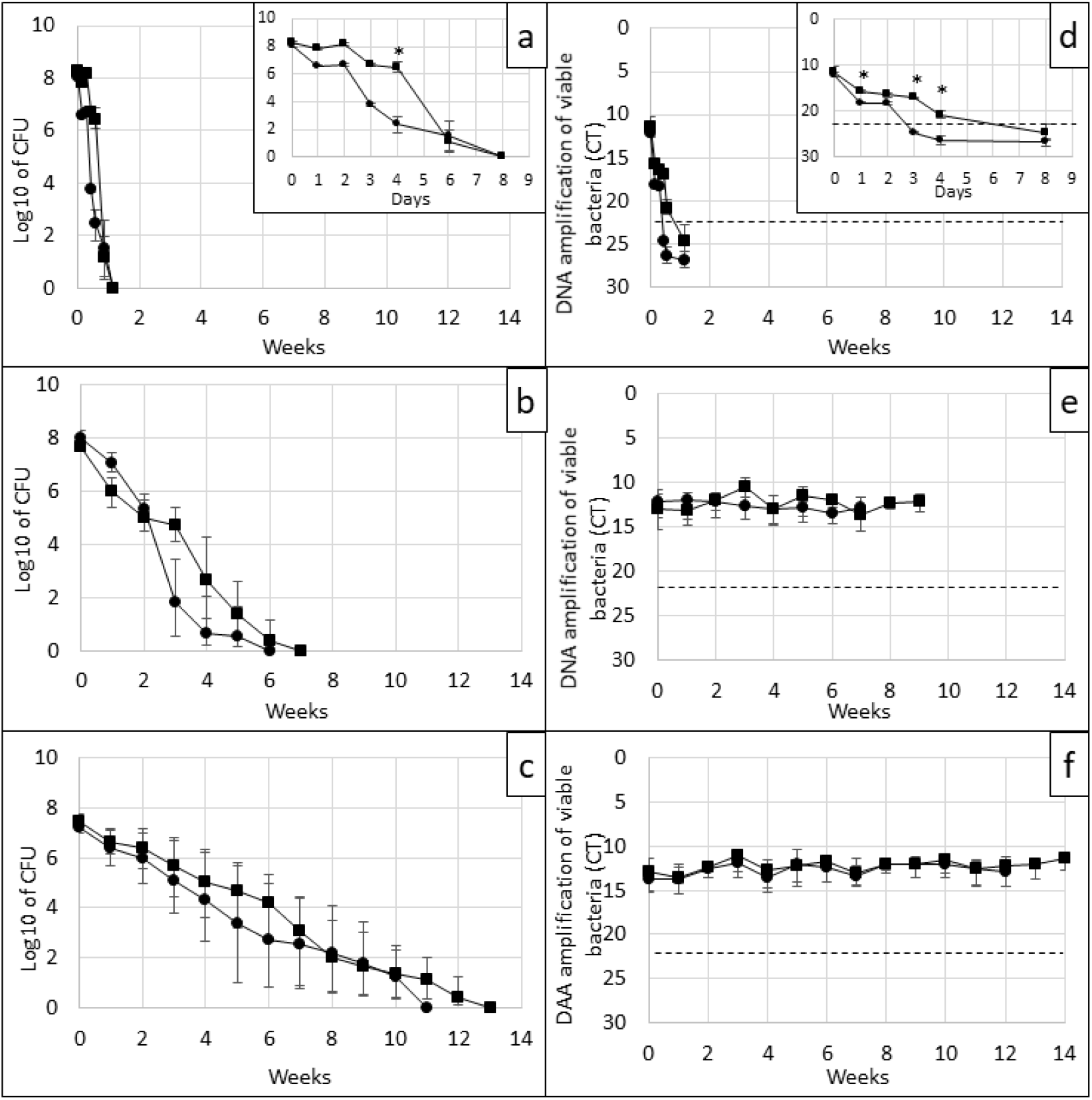
Culturability and viability of *F. tularensis* ssp. *holarctica* in nutrient-poor water. Culturability (1a-c) and viability (1d-f) of *Fth* in nutrient-poor water at respectively 37°C (1a,d), 18°C (1b,e) and 4°C (1c,f). Culturability was measured by CFU counts after serial dilutions and spreading on chocolate agar plates. Viability was evaluated by amplification of DNA after PMAxx™ Dye treatment. Black circle: nutrient-poor water with 0 g/L NaCl; black square: nutrient-poor water with 10 g/L NaCl.; Dotted line: mean Ct of all the controls performed on dead populations at each sampling points. The results are expressed as the average of three biological replicates. Data were analyzed by student t-test. * p value <0,05 between samples with and without NaCl.

### F. tularensis ssp. holarctica switched to VBNC state at low temperature in nutrient-poor water

The virulent clinical strain of *Fth* biovar I did not survive for more than eight days at 37°C in nutrient-poor water. In this microcosm, viability was correlated with culturability. qPCR-PMA Ct value increased from 12.1±0.5 to 25.8 ±0.2 after eight days in nutrient-poor water without NaCl showing a strong reduction of viable bacteria in this microcosm. In comparison, for each condition the Ct value of the controls with dead bacteria, i.e., lysed and PMAxx™ Dye treated bacteria, was 22.7±3.9. On the opposite, while the culturability declined, almost all bacteria remained alive during the eight-week study at 18°C and the 14 weeks study at 4°C (Figures 1e and 1f). The Ct value of viable bacteria stayed stable at 12.5 ±1.3 for all four conditions during the whole experiment and for more than two weeks after the loss of culturability (Water at 4°C without NaCl, Ct range: 11.8-13.8. Water at 4°C with NaCl, Ct range: 11-13.5. Water at 18°C without NaCl, Ct range: 12-13.6. Water at 18°C with NaCl, Ct range: 10.4-13.7). Two replicates were kept in the water for 24 months and tested again. Interestingly, the Ct of PMA-qPCR remained unchanged (13.2 and 14.1). Thus, we observed that roughly the full initial bacterial inoculum switched at low temperature to viable but non-culturable state corresponding to the definition of transition into VBNC state. It is interesting to note that the temperatures of 4°C and 18°C differentially affected culturability but not viability. Viability of the bacteria in the microcosms at 4 and 18°C was confirmed by the Live/Dead® BacLight™ assay (Figure S1).

The addition of 10g/L NaCl conferred a slight transient survival advantage to the bacteria at 37°C. On the fourth day, there was a 4-log difference (p-value = 0.0005) between the two conditions but qPCR-PMA Ct value also increased from 11.4±1.1 to 22.6 ±0.4 showing that all the bacteria were dead in eight days in both conditions (Figure 1d). The addition of salt to the microcosm did not significantly affect the culturability and viability of the bacteria at 4°C and 18°C (p-value > 0,05 for each time points, Figures 1b,c,e,f).

To visualize bacterial morphology after transition into VBNC state, fresh bacteria suspended one hour in water and VBNC bacteria sampled five months after the loss of culturability were labeled with an anti-*F. tularensis* LPS antibody and observed with oil immersion objective 100X. After the loss of culturability the anti-LPS antibody was still able to bind to the LPS of *Fth* strain and microscopic examination suggested a reduced length of VBNC bacteria (Figure S2). Modification of the size of the bacteria was confirmed by Syto9 staining and image analysis that showed that VBNC bacteria were smaller than culturable *Fth* with respectively an area of 0.31 ±0.19 μm^2^ and 0.47 ±0.27 μm^2^; a length of 0.62 ±0.22 μm and 0.76 ±0.26 μm; a perimeter of 1.85±0.63 μm and 2.29±0.75 μm (p-value <0.0001 for each parameters). However, circularity was not statistically different (0.97±0.03 for both culturable and VBNC *Fth* samples; p-value = 0.47) (Figure S3).

### High water temperatures inactivated F. tularensis ssp. holarctica VBNC bacteria

After their transition into the VBNC state, the viability of the bacteria was still dependent on the temperature of the water. Several months after the loss of culturability, when VBNC bacteria were moved from 4°C to 18°C and vice versa, the temperature change did not influence the viability of the bacteria during after 14 days of incubation (4°C to 18°C: Ct value from 13.8±0.8 to 14.3±2.5; 18°C to 4°C: Ct value from 12.7±1.1 to 15.3 ±3.7). However, when the temperature was shifted to 30°C, the viability of VBNC bacteria significantly declined in 14 days (4°C to 30°C: Ct value from 13.8±0.8 to 20±1.5; 18°C to 30°C: Ct value from 12.7±1.1 to 20±6) (p-value < 0.05). Moreover, when placed at 37°C, viability of VBNC bacteria declined in 14 days under the threshold corresponding to dead bacteria only (4°C to 37°C: Ct value from 13.8±0.8 to 23.8±2.5; 18°C to 37°C: Ct value from 12.7±1.1 to 26.3±3.3) (p-value <0.05) (Figure 2).

**Figure 2:**
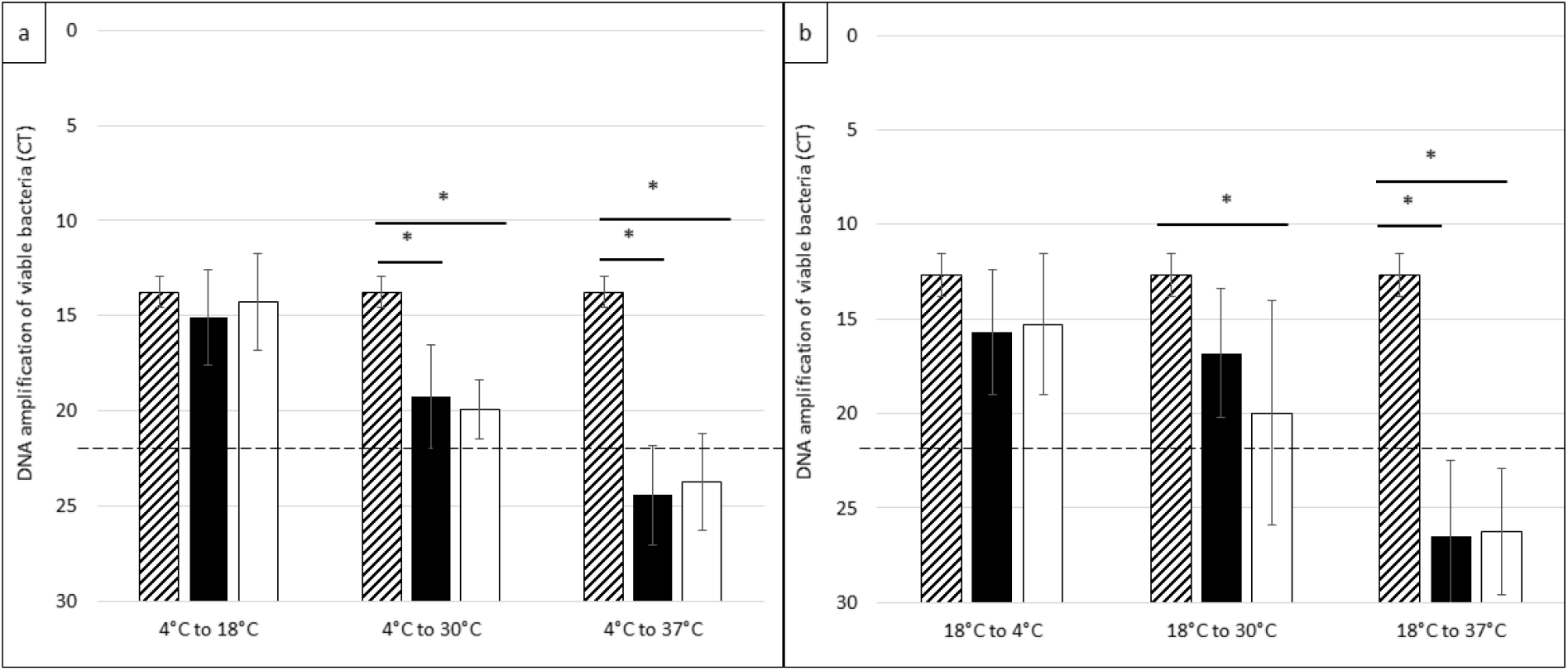
Viability of VBNC *F. tularensis* ssp. *holarctica* in water after a temperature change. Several months after the loss of culturability of *Fth* in water, 5 mL of microcosm at 4°C were transferred at 18°C, 30°C and 37°C (3a) and 5 mL of microcosm at 18°C were transferred at 4°C, 30°C and 37°C (3b). After 7 and 14 days, the viability was evaluated by qPCR after PMAxx™ Dye treatment. Dash bars: Ct at day 0, black bars: Ct at day 7, white bars: Ct at day 14, dotted line: Ct of the control corresponding to average of a dead population. The results are expressed as the average of three biological replicates. Data were analyzed by one-way ANOVA with pairwise t-tests using FDR correction. * p-value < 0.05.

### Virulent F. tularensis ssp. holarctica strain was able to form biofilm in water

Optical density at 570 nm of the water flasks containing fresh bacteria after crystal violet staining was 0.14±0.01 while optical density of the flasks containing the *Fth* VBNC bacteria were increased twofold: 0.27±0.01 for VBNC bacteria after one year at 4°C (p-value = 0.019) and 0.3±0.01 (p-value < 0.0001) for VBNC bacteria after six months at 18°C. On the opposite, optical density of the flasks containing dead *Fth* bacteria (4 months at 37°C) was 0.15±0.03 showing no significant biofilm production compared to fresh bacteria (p-value = 0.7) (Figure 3). Microscopic observation of the stained flasks showed small Gram-negative coccobacilli embedded and surrounded by a structure resembling a biofilm (Figure S4).

**Figure 3:**
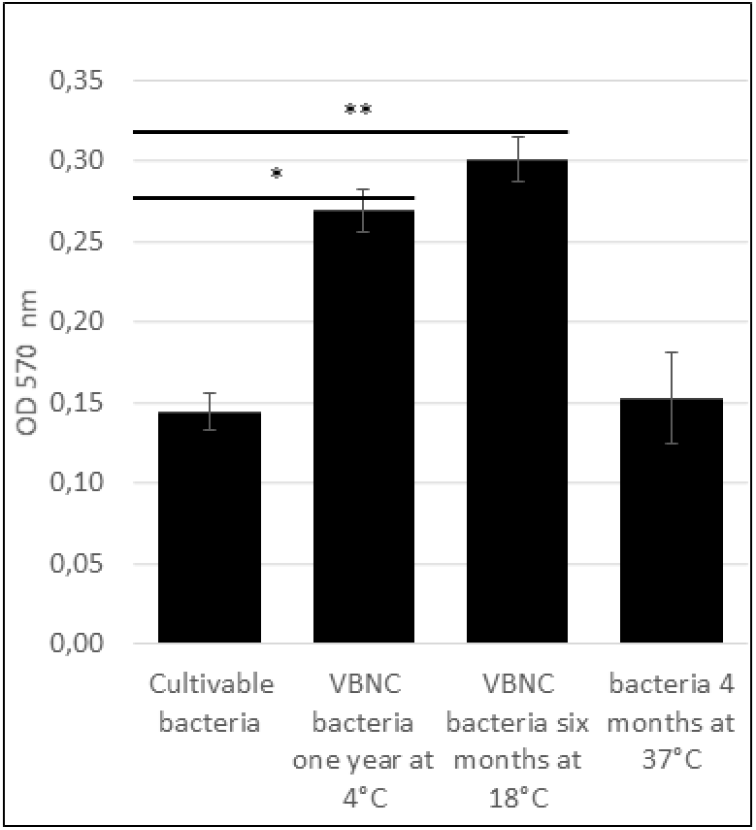
Quantitative measurement of biofilm formation of *F. tularensis* ssp. *holarctica* in VBNC state. *Fth* bacteria were incubated in nutrient-poor water for one hour for cultivable bacteria, one year at 4°C and six months at 18°C for VBNC and for four months at 37°C for non-persistent bacteria. Biofilm biomass was estimated by absorbance at 570 nm of crystal violet assay. The results are expressed as the average of three biological replicates. Data were analyzed by student t-test, * p-value <0.05 ** p-value < 0.01.

## Discussion

Although the presence and potential survival of *Fth* in the aquatic environment have been identified in several studies [25–29], the mechanisms of its persistence and its precise environmental reservoir remain unclear. According to current descriptions of the aquatic cycle of *Fth*, aquatic environments may be initially contaminated by *F. tularensis* through dead animals or excrements of infected animals [3]. However, how these bacteria can persist for weeks or even years within these environments remains to be elucidated. Following recent work showing extended culturability of *Fth* at 4°C in water, we hypothesized that in environmental water, this bacterium might also survive in a dormancy form such as the VBNC state. This hypothesis would help the bacteria to survive in hostile environments, as described for several other Gram-negative bacteria, thus limiting nutrient starvation and competition with other microorganisms [20,22,30,31].

We observed that a clinical strain of *Fth* remained culturable for more than 11 weeks of incubation in nutrient-poor water at 4°C; more than one month at 18°C but only one week at 37°C, consistent with previously published data on the culturability of *Fth* FSC200 and LVS [4,19–21]. However, we show here that culturability is not representative of the viability of *Fth* strains since the bacterium may switch to the VBNC state under conditions that remain to be fully characterized. Indeed, our results showed prolonged survival in nutrient-poor water at 4°C and 18°C of a virulent *Fth* biovar I strain long after the bacterium had lost its ability to grow on an agar plate. Our main approach assessing bacterial survival is based on qPCR amplification of DNA from bacteria preincubated with PMAxx™ Dye widely used to detect and determine the viability of human pathogens [32]. As PMAxx™ Dye does not pass through intact bacterial membranes, it cannot bind to the DNA of living bacteria although binding to the DNA of dead bacteria and extracellular DNA is possible. While the amount of DNA from living bacteria decreased similarly to culturability at 37°C, it remained remarkably stable over time at 4°C and 18°C during the whole experiment matching the definition of bacterial switch into a VBNC state as the majority of initial bacteria remained viable despite the loss of culturability and results were confirmed by Live/Dead® BacLight™ assay [33]. Morphological analysis showed that VBNC *Fth* bacteria are smaller as they have a reduced length, perimeter and area compared to the culturable forms.

Like the seeds of plants, VBNC forms allow preserving the genetic heritage of bacteria in unfavorable conditions [34]. *Fth* bacteria could then remain viable for a very long time as VBNC bacteria in aquatic environments without the need for a host. When more favorable conditions return, VBNC bacteria revert to their vegetative state, usually recovering their culturability and virulence. Reversion after switch into VBNC state remains to be demonstrated for *Fth* in further studies.

Bacteria evolve to a VBNC state to withstand environmentally induced stresses. In our experiments, incubation of *Fth* in water at 37°C was the most deleterious environmental condition. It did not induce a transition to the VBNC state since bacterial mortality correlated with loss of culturability. Therefore, it appears that conditions that are too harmful to *Fth* and associated with a loss of their culturability in one week do not allow the development of VBNC bacteria. The most favorable conditions for *Fth* survival are close to environmental conditions in tularemia endemic areas, i.e., areas of water temperatures ranging from 4 to 20°C between winter and summer periods [4,28]. Importantly after the switch into VBNC state, *Fth* viability was still dependent on the temperature of the water. Over 30°C, the viability of VBNC bacteria declined, and was completely abolished after seven days at 37°C. Thus, the temperature tipping point no longer supporting the transition of *Fth* to the VBNC state is between 18°C and 30°C. Our results support the aquatic environmental distribution of *Fth* in Northern regions where water temperature may not often exceed the temperature limit killing bacteria in a VBNC state. The inter-tropical region, with higher water temperature, could therefore represents a physical limitation to spreading towards the southern hemisphere. The seasonality could also have a significant role in maintaining this environmental reservoir since the bacterial persistence is better in freshwater.

One other mode of persistence of bacteria in aquatic environments is biofilm formation, as observed for many bacteria like *Legionella pneumophila* [35]. Experimental studies have shown that environmental species of *Francisella* can form biofilms *in vitro* [36]. *F. novicida* starts biofilm formation after two hours and can be evidenced by crystal violet staining after 24h [36]. In our study, we observed thin biofilm formation at the bottom of the flasks after six or 12 months of incubation of the microcosms at 18°C or 4°C but no biofilm formation after four months at 37°C. The biofilm was very fragile and therefore difficult to manipulate for observation. Biofilm formation of *Fth* strains may be a slow process requiring the viability of the bacteria for more than one week. The absence of biofilm formation of *F. tularensis* strains observed in the study of Golovliov *et al*. may be related to experimental conditions in axenic media not mimicking the natural aquatic environment [4]. Our study is closer to environmental conditions, although it did not contain other competitive bacteria or predatory microorganism, because the *Fth* strain was incubated in a large volume of filtered French lake water. In our experiment, VBNC bacteria seemed to be embedded in a biofilm matrix, as previously shown for other pathogens (e.g., *Legionella pneumophila* and *Listeria monocytogenes*) [30,37]. In 2016, Flemming *et al*. described biofilms as a “reservoir of VBNC bacteria,” especially in the starvation zones of the biofilm [35]. The biofilm and VBNC states play an essential role in the persistence of bacteria. Both allow the bacteria to survive in hostile environments while many pathogens lose their virulence properties after their switch into a VBNC state [31].

The persistence of *Fth* in aquatic environments in a VBNC state questions our capacity to detect and fight this bacterium in this specific reservoir. VBNC state may be a way of long-term bacterial persistence of *Fth* that cannot be detected by conventional culture-based techniques. The VBNC formation process likely explains that detection of *F. tularensis* in the aquatic environment has been obtained by species-specific molecular methods but very rarely by culture techniques [5]. In case of accidental or intentional dispersal of *F. tularensis*, the bacterium may thus survive for many months in water environments although undetected by culture methods. Identification of reactivation factors from the VBNC state into a more virulent and culturable state will have to be addressed in further experiments. It would help prevent and control waterborne sporadic and outbreak tularemia cases.

The impact of temperature may also have some effects on tularemia diagnosis in humans. This bacterium is usually grown at 37°C from clinical samples in only 10% of tularemia patients. In the light of this work, this temperature may not be optimal for isolating this bacterium from patients, animal samples, or environmental samples or even for bacterial counts after growth on an agar medium. Potential switch into VBNC state *in vivo* in infected tissues (especially lymph nodes) could also partly explain the failure to isolate this bacterium and may impact therapeutic outcome as VBNC bacteria usually exhibit increased antibiotic resistance because of a reduced metabolism while biofilms also increase resistance to antibiotics [31,38]. These findings may have implications in treatment failures observed in 20 to 30% of patients, especially when the diagnosis is delayed, which could allow the bacteria to switch to a VBNC state *in vivo*.

Finally, all these data about the environmental survival of *Fth* in water at low temperature brings new important features allowing updating the aquatic cycle of this bacterium and proposing new hypotheses (Figure 4). Indeed, many other *Francisella* species are aquatic bacteria, making several parts of this aquatic cycle questionable [5]. What if *Fth* had rather evolved to adapt to an aquatic niche yet poorly characterized so far while becoming infectious for various mammal species, which are usually dead ends for the bacteria because it often kills its hosts [3]? Indeed, this bacterium has been identified in many aquatic areas, including the sea water, rivers, ponds [25–28,39,40], which might suggest that the primary reservoir of the subspecies *holarctica* could rather be the aquatic environment itself. This hypothesis could explain why a specific reservoir within the environment has not been identified despite decades of research. Some aquatic environments may thus represent the largest reservoir of *Fth* with the implication of aquatic rodents to maintain the cycle through bacterial inoculum amplification. Animals and humans may thus be infected directly from this environmental reservoir through water consumption or aerosols, explaining some sporadic human respiratory contaminations after outdoor activities. Aquatic environments could act as a primary source of human and animal infections or mosquito larvae contamination after reactivation of the VBNC state into virulent bacteria upon particular environmental conditions. Further studies will be necessary to determine: 1/ if *Fth* VBNC bacteria are also virulent and able to infect animals and mosquito larvae who are at the interface between the aquatic and the terrestrial cycle; 2/ to identify reactivation factors from the VBNC state towards the culturable and virulent state; 3/ to study interaction of VBNC *Fth* bacteria within biofilms and with amoeba.

**Figure 4:**
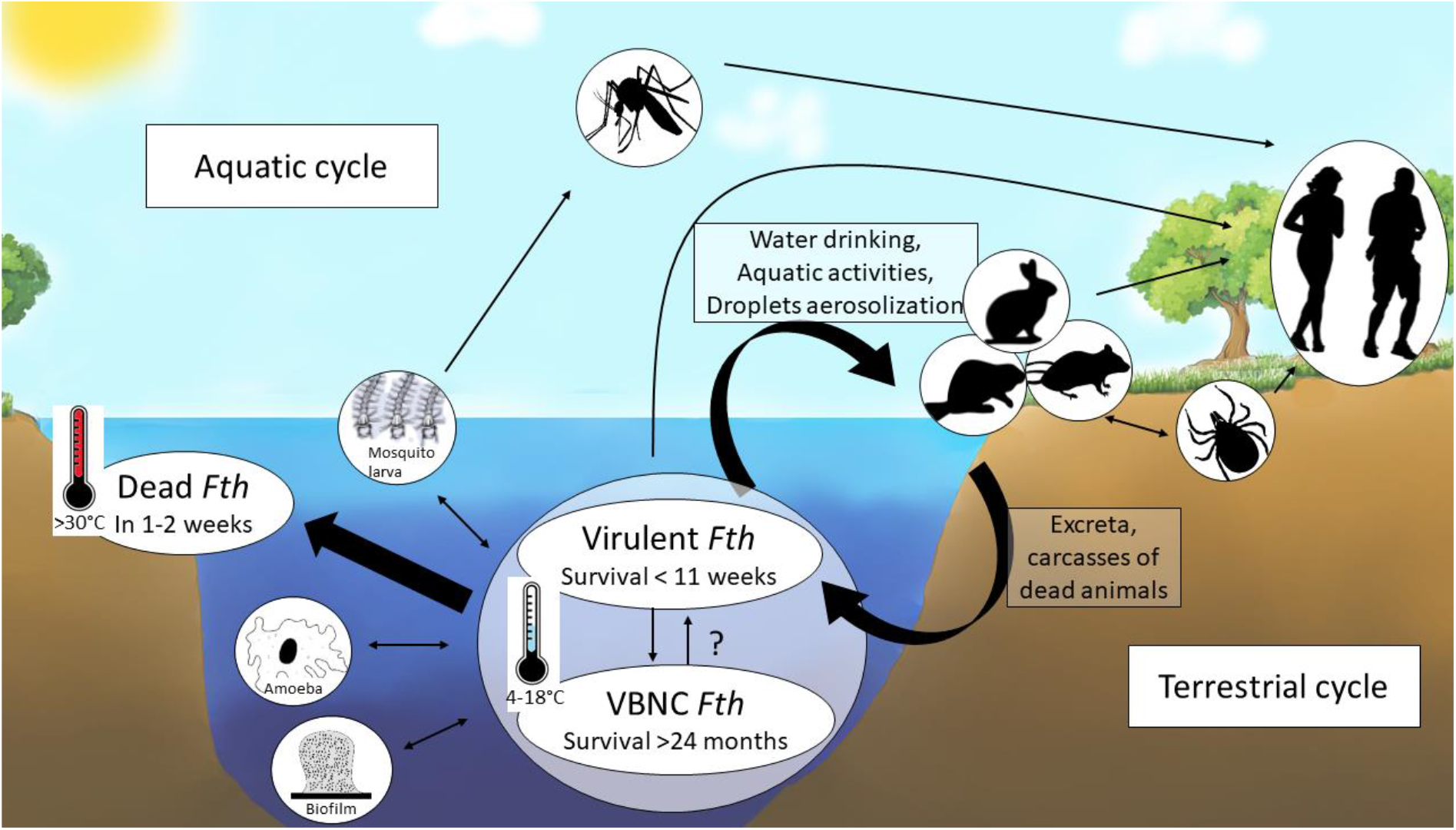
The hidden aquatic reservoir of *F. tularensis* ssp. *holarctica*? We updated the current knowledge about the aquatic cycle of *Fth* according to the results of this work and proposed hypotheses that emerged from our observations. We showed that the survival of *Fth* in aquatic environments is driven by water temperature and transition into a VBNC state. While *Fth* culturability is prolonged in water at low temperatures (4-18°C), these low temperatures actually also allow the survival of the bacteria for months or years after transition into a VBNC state. On the opposite high temperatures (> 30°C) are associated to complete loss of culturability and loss of viability of the bacteria, even if the bacteria has already switched into the VBNC state at lower temperatures. Thus, mammals or accidentally human may be contaminated from this long-term aquatic reservoir by water drinking, direct contact or by inhalation of contaminated droplets that could explain several respiratory tularemia cases related to environmental exposure only. When infected, wild animals can amplify the bacterial inoculum within the same aquatic environment or disperse the bacteria in other environments with their carcasses and feces and may contaminate other animals or exceptionally humans as described in the terrestrial cycle of the bacteria.

In conclusion, our study demonstrated the extended persistence of a virulent strain of *Fth* in water up to 24 months through the formation of VBNC bacteria and thin biofilms. Water temperature appears as a major factor for bacterial survival in aquatic environments. It affects the culturability of the bacteria, the switch toward the VBNC state, and the viability of VBNC cells. Our findings reinforce the hypothesis of a long-term environmental aquatic reservoir of this pathogen.

## Acknowledgment

We thank the company Abiolab Asposan for the chemical analysis of the water. We thank Ludovic Sansoni for his help on the figure of aquatic cycle.

