## supplemental table and figure for "Aquatic long-term persistence of *Francisella tularensis* ssp. *holarctica* is driven by water temperature and transition to a viable but non-culturable state" (from another version of this preprint: version 2, 2022-02-21)

**Table S1: Physico-chemical analysis of environmental water**

<u>Material & Methods</u>

The water used for the models was taken in winter from a French lake. pH, calcium, nitrates, nitrites, orthophosphates, sulfates, chloride, sodium, potassium, magnesium, dissolved oxygen, hydrogen carbonates, and anionic detergents were measured (Abiolab-Asposan, Monbonnot-Saint-Martin, France) in one liter of water with *Fth* at a concentration of $10^8$ bacteria / mL incubated at 18°C. Two weeks after the loss of bacterial culturability, samples were filtered through 0,22µm filters and sent for physicochemical analysis. Another 1 liter of water was not inoculated with bacteria but processed and analyzed similarly.

<u>Results</u>

pH, calcium, nitrates, nitrites, orthophosphates, sulfates, chloride, sodium, potassium, magnesium, dissolved oxygen, hydrogen carbonates, and anionic detergents were measured before and after incubation of bacteria. These measurements confirmed that water used without NaCl addition contained 0 g/L NaCl. The bacteria did not change any parameters measured during the eight weeks of incubation. Microcosm were filled with this environmental water, considered as low nutrient water as it contains less nutrient than usual culture media for *Fth* but more than saline solution.

<u>Caption</u>

Physico-chemical analysis of environmental water

| | Without bacteria incubation | With bacteria incubated 8 weeks at 18°C |
|---|---|---|
| pH | Neutral | Neutral |
| Calcium (Ca) | 50 mg/L | 51 mg/L |
| Nitrates (NO3) | <0.5 mg/L | <0.5 mg/L |
| Nitrites (NO2) | <0.02 mg/L | <0.02 mg/L |
| Orthophosphates (PO4) | 2.98 mg/L | 2.99 mg/L |
| Sulphates (SO4) | 29 mg/L | 30 mg/L |
| Total chlorine | <0.02 mgCl2/L | <0.02 mgCl2/L |
| Sodium (Na) | 350 mg/L | 360 mg/L |
| Potassium (K) | 11 mg/L | 11 mg/L |
| Magnesium (Mg) | 14 mg/L | 14 mg/L |
| Dissolved oxygen | 8.3 mg/L | 8.2 mg/L |
| Hydrogen carbonates (HCO3) | 212 mg/L | 218 mg/L |
| Anionic detergents | <0.1 mg/L | <0.1 mg/L |

**Figure S1: Live/Dead® BacLight™ labelling**

Material & Methods

The Live/Dead® BacLight™ assay distinguishes dead bacteria with mixt Syto9 and Propidium Iodide (PI). All bacteria were labeled in green by Syto9 and dead bacteria were labeled in red by PI. Briefly, 130 µL of culture were diluted and incubated with a mixture containing 0.2 µL of PI and 0.2 µL of Syto9. After 15 min of incubation in the dark, bacteria were observed using a fluorescence microscope with oil immersion objective 100Xa/1.25 (Nikon).

Results

Six months after their inoculation in microcosm, only a few numbers of bacteria incubated at 4°C and 18°C and visualized by Syto9 were stained by IP. On the contrary, one month after their inoculation in microcosm, all bacteria incubated at 37°C were marked by Syto9 and IP, demonstrating that there were no more living bacteria in this experimental condition.

Caption

Fluorescent live/dead staining of *F. tularensis* subsp. *holarctica*. Bacteria were incubated in nutrient-poor water and treated by Live/Dead® BacLight™ kit after six months incubation at 4°C or 18°C, and after one month at 37°C.

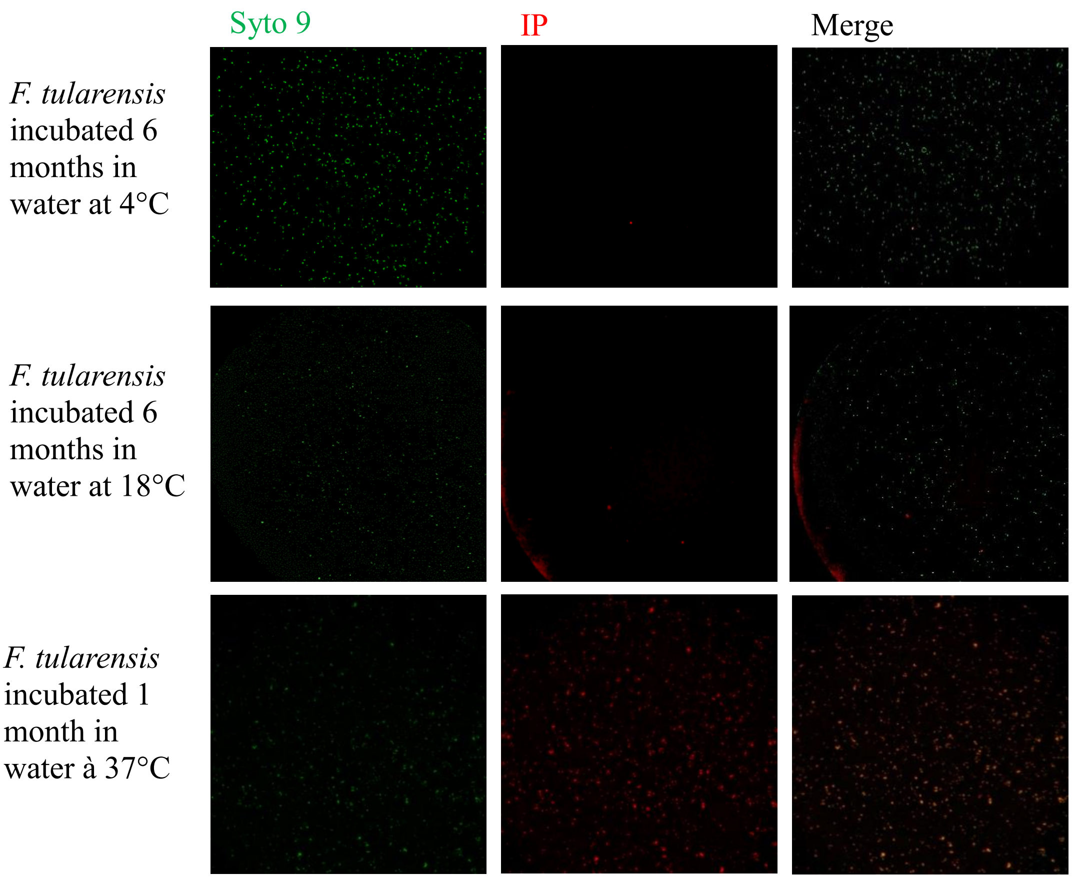

**Figure S2: Fluorescent labeling of bacteria**

Material & Methods

Imaging was performed on culturable and non-culturable bacteria to observe the shape of bacteria before and after their switch into VBNC state, respectively. VBNC bacteria were induced in water at 18°C, and observed five months after the loss of culturability. Culturable bacteria corresponded to fresh bacteria grown on PVX agar plate and suspended in water at 18°C one hour. 1 ml of each sample was added to a well with a coverslip. Bacteria were fixed on slides with PFA 4% for 10 min. The slides were then saturated with BSA 3% for 30 min and were covered with a primary anti-*F. tularensis* LPS antibody (ThermoFisher Scientific, Waltham, US) diluted at 1/100, and incubated for 1 hour at room temperature. After washing, the slide was covered with a secondary anti-mouse antibody coupled to the Alexa fluor 488 (ThermoFisher Scientific, Waltham, US) at 1/1000 dilution and incubated for 1 hour at room temperature. Microscopic observation was realized using a scanning confocal microscope (Zeiss LSM710) with oil immersion objective 100X.

Caption

Fluorescent labeling of *F. tularensis* ssp. *holarctica* in culturable form (2a) and VBNC form (2b). Bacteria-LPS were labeled with a primary anti-*F. tularensis* LPS antibody (ThermoFisher Scientific, Waltham, US) and a secondary anti-mouse antibody coupled to the Alexa fluor 488 (ThermoFisher Scientific, Waltham, US). Microscopic observation was realized using a scanning confocal microscope (Zeiss LSM710) with oil immersion objective 100X.

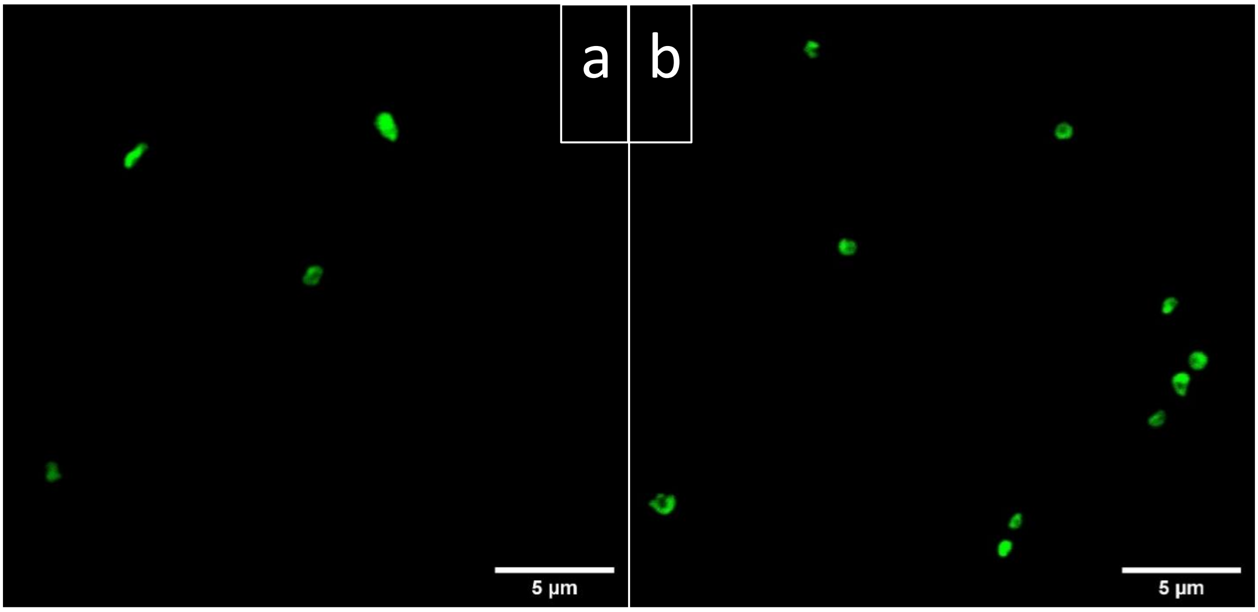

**Figure S3: Bacterial morphology**

Material & Methods

To compare the shape and the length of bacteria, pictures of bacteria labeled with Syto9 were analyzed by Image J software and Microbe J plugging[1]. Bacterial morphology was described by parameters: area (0.1-1.2µm); length (0.2-1.4 µm); width (0-1.4 µm); circularity (0.3-max µm). Parameters were calculated for 1055 bacteria in each sample.

Results

Modification of the size of the bacteria was confirmed by Syto9 staining and image analysis that showed that VBNC *Fth* were smaller than culturable. *Fth* with respectively an area of 0.31 ±0.19 µm² and 0.47 ±0.27 µm²; a length of 0.62 ±0.22 µm and 0.76 ±0.26 µm; a perimeter of 1.85 ±0.63 µm and 2.29 ±0.75 µm (p-value <0.0001 for each parameters). However, VBNC bacteria kept a similar round shape with a circularity of 0.97 ±0.03 for both culturable and VBNC *Fth* sample (p-value = 0.47).

Caption

Bacterial morphology of culturable *Fth* (white bars) and VBNC *Fth* (black bars) in µm. Data were analyzed by student t-test ** p-value < 0.005.

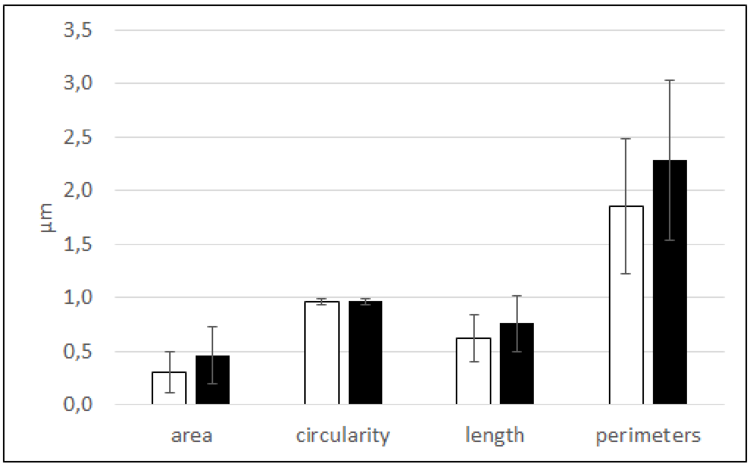


[1] A. Ducret, E.M. Quardokus and Y.V. Brun, *MicrobeJ, a tool for high throughput bacterial cell detection and quantitative analysis*, Nat. Microbiol. 1 (2016), pp. 16077.

**Figure S4: Microscopic examination of biofilm**

Material & Methods

Biofilm labeling was performed on bacterial suspension evolved in nutrient-poor water in T75 culture flasks at 18°C for six months. The culture supernatant was aspirated, and the flask was washed three times with PBS. 5 mL of Cristal violet (0,2% w/v) were added and incubated for 15 minutes. After three washing, the flask was opened, biofilm was transferred on a coverslip, placed on a glass slide, and fixed by Eukitt®. Biofilm was observed by an optical microscope (oil immersion objective 100X).

Caption

*F. tularensis* ssp. *holarctica* biofilm containing VBNC bacteria.
Bacteria were incubated in nutrient-poor water at 18°C in a culture flask; after six months, flasks were washed, and biofilms were stained with crystal violet. Biofilms were observed by optic microscope (oil immersion objective 100X).

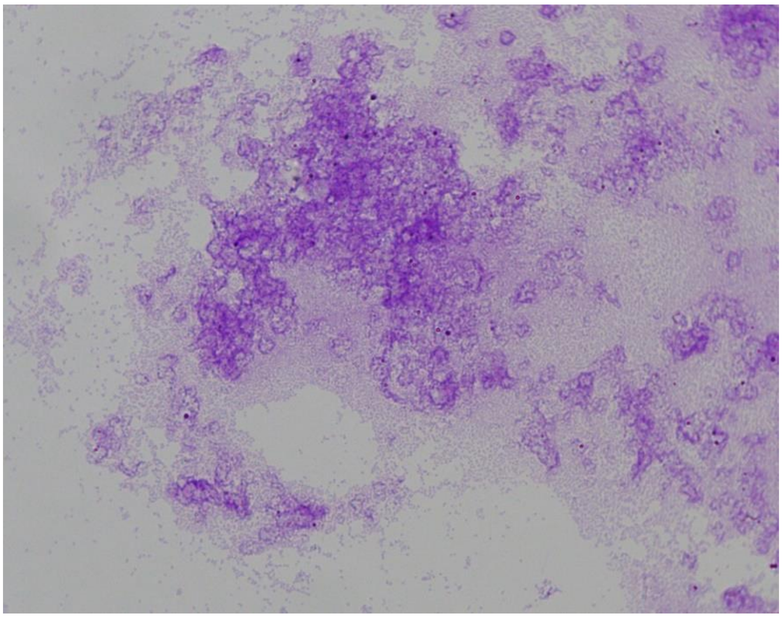
